# Brain *de novo* transcriptome assembly of a toad species showing polymorphic anti-predatory behavior

**DOI:** 10.1101/2022.08.03.502642

**Authors:** Andrea Chiocchio, Pietro Libro, Giuseppe Martino, Roberta Bisconti, Tiziana Castrignanò, Daniele Canestrelli

## Abstract

Understanding the genomic underpinnings of antipredatory behaviors is a hot topic in eco-evolutionary research. Yellow-bellied toad of the genus *Bombina* are textbook examples of the deimatic display, a time-structured behavior aimed at startling predators. Here, we generated the first *de novo* brain transcriptome of the Apennine yellow-bellied toad *Bombina pachypus,* a species showing inter-individual variation in the deimatic display. Through Rna-Seq experiments on a set of individuals showing distinct behavioral phenotypes, we generated 316,329,573 reads, which were assembled and annotated. The high-quality assembly was confirmed by assembly validators and by aligning the contigs against the *de novo* transcriptome with a mapping percentage higher than 91.0%. The homology annotation with DIAMOND (blastx) led to 77,391 contigs annotated on Nr, Swiss Prot and TrEMBL, whereas the domain and site protein prediction made with InterProScan led to 4747 GO-annotated and 1025 KEGG-annotated contigs. The *B. pachypus* transcriptome described here will be a valuable resource for further studies on the genomic underpinnings of behavioral variation in amphibians.

## Background & Summary

Inter-individual variation in antipredatory behavior has long attracted scientific curiosity and has been investigated in a wide range of animal species, from mammals to fishes, insects and even to marine invertebrates^1^. However, we still know little about the specific molecular mechanisms underlying the origin of this variation. In fact, while some behavioral traits have been linked to epigenetic mechanisms^2^, the observation that behavior can be heritable supports a role for modulation of standing genetic variation within populations^3–4^. The recent advances in omic-sciences opens for investigating the genetic basis of behavioral trait variation^5^, and thus for understanding the genomic underpinnings of the inter-individual variation in antipredatory strategies.

Deimatism is a common anti-predatory strategy. It consists in suddenly unleashing unexpected defenses to frighten predators and to stop their attack, and it combines cryptism and aposematism in a complex and time structured antipredatory strategy^6^. Deimatic displays typically involve either chromatic and behavioral components. Inter-individual variation in warning signals have traditionally been considered maladaptive. Yet, individual variation in morphological and chromatic components have been widely reported in many organisms^7–11^. Conversely, the occurrence of polymorphism in the behavioral component of warning signals is still almost unexplored.

With this study, we aim at providing genomic resources to investigate the genetic underpinnings of inter-individual behavioral differences in warning signals. We generated the first *de novo* brain transcriptome of a species showing polymorphism in behavioral traits associated with deimatic displays, the Apennine yellow-bellied toad *Bombina pachypus*^12^. The yellow-bellied toads of the genus *Bombina* are textbook examples of unken-reflex, a deimatic behavior which consists in toads arching their body and exposing their aposematically colored ventral side. Experimental evidence has shown within-population variation in the way *B. pachypus* toads reacted to predation stimuli: about half of the toads quickly reacted with a long and intense body arching and aposematic display (i.e. the unken-reflex), while the other half of the individuals analysed did not show deimatic behavior, but rather moved away^12^. These two antipredatory strategies have been proposed to reflect the way individuals cope with environmental challenges, i.e. reactive vs proactive coping style, and arguably linked to differential sympathetic-parasympathetic activities^13^. We focused on brain transcriptome, as the brain tissues have shown differential gene expression profiles linked to distinct behavioral states in response to environmental stimuli^14–16^, also in closely related *Bombina* species^17–18^. Thus, the transcriptome analysis of the brain can reveal the ways in which distinct molecular pathways can modulate anti-predatory behaviour^19^.

The data presented in this study consist of assembled transcriptome sequences of the brain of *B. pachypus* at the adult stage. The *de novo* transcriptome has been annotated to provide a transcriptome reference for further analysis of differential gene expression profiles. Since the *B. pachypus* genome has not been sequenced so far, the transcriptome presented here will be a valuable resource for further eco-evolutionary studies on the behavioral repertoire of amphibians.

## Methods

### Sample collection and RNA preparation

We analyzed 6 adult yellow-bellied toad individuals representative of distinct behavioral profiles, i.e. prolonged unken-reflex display vs no unken-reflex display (thereafter referred as “+” and “-”, respectively). Behavioral profiles were scored as in Chiocchio et al.^12^: 3 toads showed prolonged unken-reflex (+), whereas the other 3 did not show unken-reflex (-), as reported in Table 1. Sampling procedures were approved by the Italian Ministry of Ecological Transition and the Italian National Institute for Environmental Protection and Research (ISPRA; permit number: 20824, 18-03-2020). After dissection, brain tissue was immediately stored in RNAprotect Tissue Reagent (Quiagen) until RNA extraction. RNA extractions were performed using the RNeasy Plus Kit (Quiagen), according to the manufacturer’ instructions. RNA quality and concentration were assessed by means of both a spectrophotometer and a Bioanalyzer (Agilent Cary60 UV-vis and Agilent 2100, respectively - Agilent Technologies, Santa Clara, USA).

**Table 1.** Summary of the 6 libraries deposited in the Sequence Read Archive (SRA) of NCBI, in terms of number of raw and trimmed reads per sample.

| Bioproject | Run ID | Phenotypes | Raw sequences | Filtered sequences (% of the total reads) |
| --- | --- | --- | --- | --- |
| PRJNA764013 | SRR15927734 | not unken-reflex (-) | 52476660 | 49488968 (94.31 %) |
| PRJNA764013 | SRR15927733 | not unken-reflex (-) | 52548719 | 49298011 (93.81 %) |
| PRJNA764013 | SRR15927731 | unken-reflex (+) | 53007498 | 49675835 (93.71 %) |
| PRJNA764013 | SRR15927732 | not unken-reflex (-) | 54267911 | 50813920 (93.64 %) |
| PRJNA764013 | SRR15927730 | unken-reflex (+) | 51029133 | 48227054 (94.51 %) |
| PRJNA764013 | SRR15927729 | unken-reflex (+) | 52999652 | 49850617 (94.06 %) |

### Library preparation and sequencing

Library preparation and RNA sequencing were performed by NOVOGENE (UK) COMPANY LIMITED using Illumina NovaSeq platform. Library construction was carried out using the NEBNext^®^ Ultra ™ RNA Library Prep Kit for Illumina^®^, following manufacturer instructions. Briefly, after the quality control check, the mRNA sample was isolated from the total RNA by using magnetic beads made of oligos d(T)25 (i.e. polyA-tail mRNA enrichment). Subsequently, mRNA was randomly fragmented, and a cDNA synthesis step proceeded using random hexamers and the reverse transcriptase enzyme. Once the synthesis of the first chain has finished, the second chain was synthesized with the addition of the Illumina buffer, dNTPs, RNase H and polymerase I of *E.coli,* by means of the Nick translation method. Then, the resulting products went through purification, repair, A-tailing and adapter ligation. Fragments of the appropriate size were enriched by PCR, the indexed P5 and P7 primers were introduced, and the final products were purified. Finally, the Illumina Novaseq6000 sequencing system was used to sequence the libraries, through a paired-end 150bp (PE150) strategy. We obtained on average 52.7 million reads for each library. The sequencing data are available at the NCBI Sequence Read Archive (project ID PRJNA764013).

### Pre-assembly processing stage

A total of 316,329,573 pairs of reads was generated by Illumina sequencing. All of them went to a cleaning analytic step. The quality of the raw reads was assessed with the FastQC 0.11.5 tool (http://www.bioinformatics.bbsrc.ac.uk/projects/fastqc), in order to estimate the RNAseq quality profiles. The quality estimators were generated for both the raw and trimmed data. The quality assessment metrics for trimmed data were aggregated across all samples into a single report for a summary visualization with MultiQC software tool^20^ v.1.9 (see Figure 1). To remove low quality bases and adapter sequences, raw reads were also analyzed through a quality trimming step with Trimmomatic^21^, v.0.39 (setting the option SLIDINGWINDOW: 4: 15, MINLEN: 36, and HEADCROP: 13). All the unpaired reads were discarded. After the cleaning step and removal of low-quality reads, 297,354,405 clean reads (i.e. 94% of raw reads) were maintained for building the *de novo* transcriptome assembly (see Table 1).

**Figure 1.**
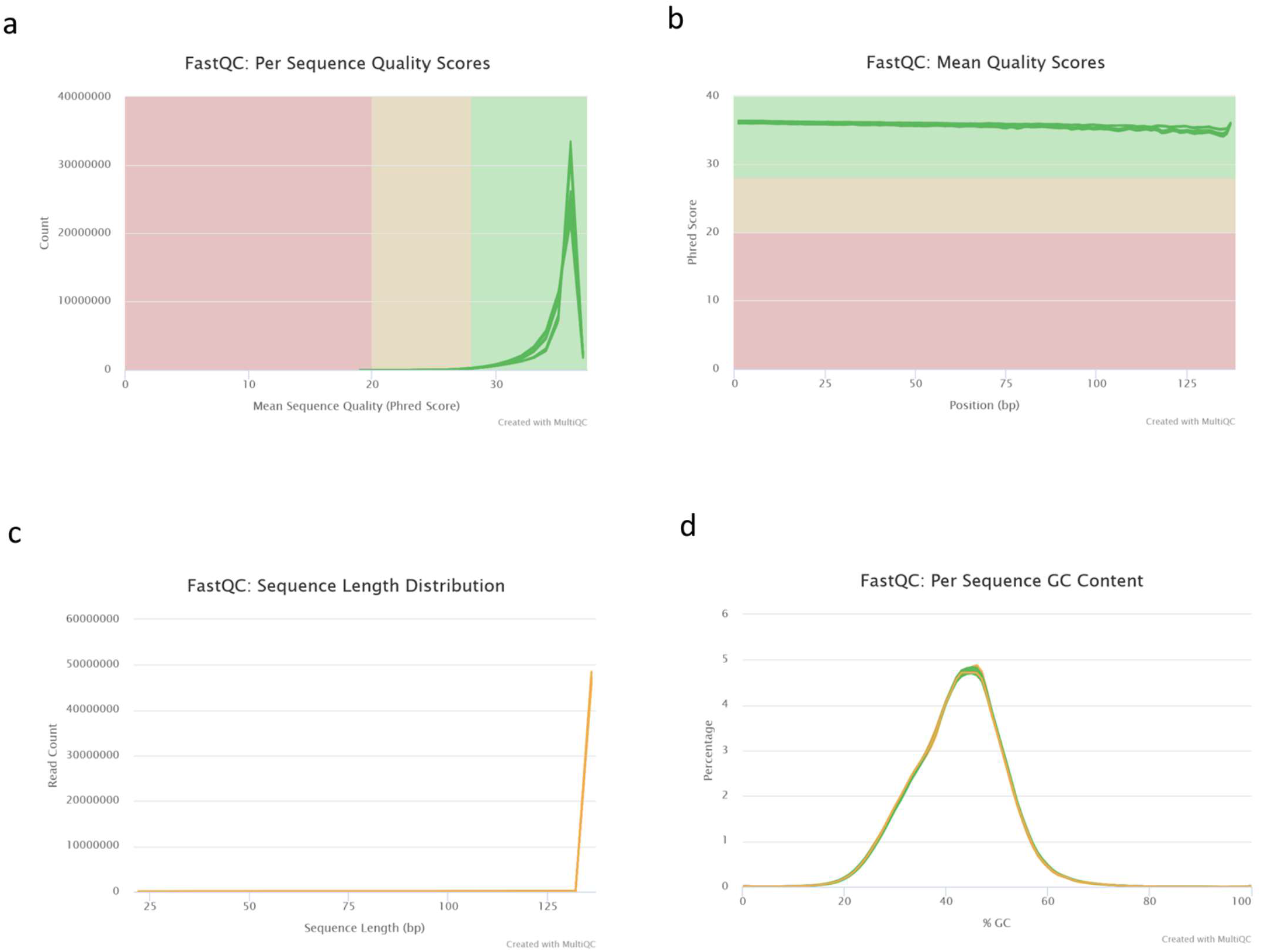
The cleaned reads from all samples were assessed with FastQC and visualized with MultiQC. (a) Read count distribution for mean sequence quality. (b) Mean quality scores distribution. (c) Read length distribution. (d) Per Sequence GC Content

### *De novo* transcriptome assembly and quality assessment

As there is no reference genome for *B. pachypus,* we performed a *de novo* transcriptome assembly procedure. The workflow of the bioinformatic pipelines is shown in Figure 2. All the described bioinformatics analyses were performed on the high-performance computing systems provided by ELIXIR-IT HPC@CINECA^22^.

**Figure 2.**
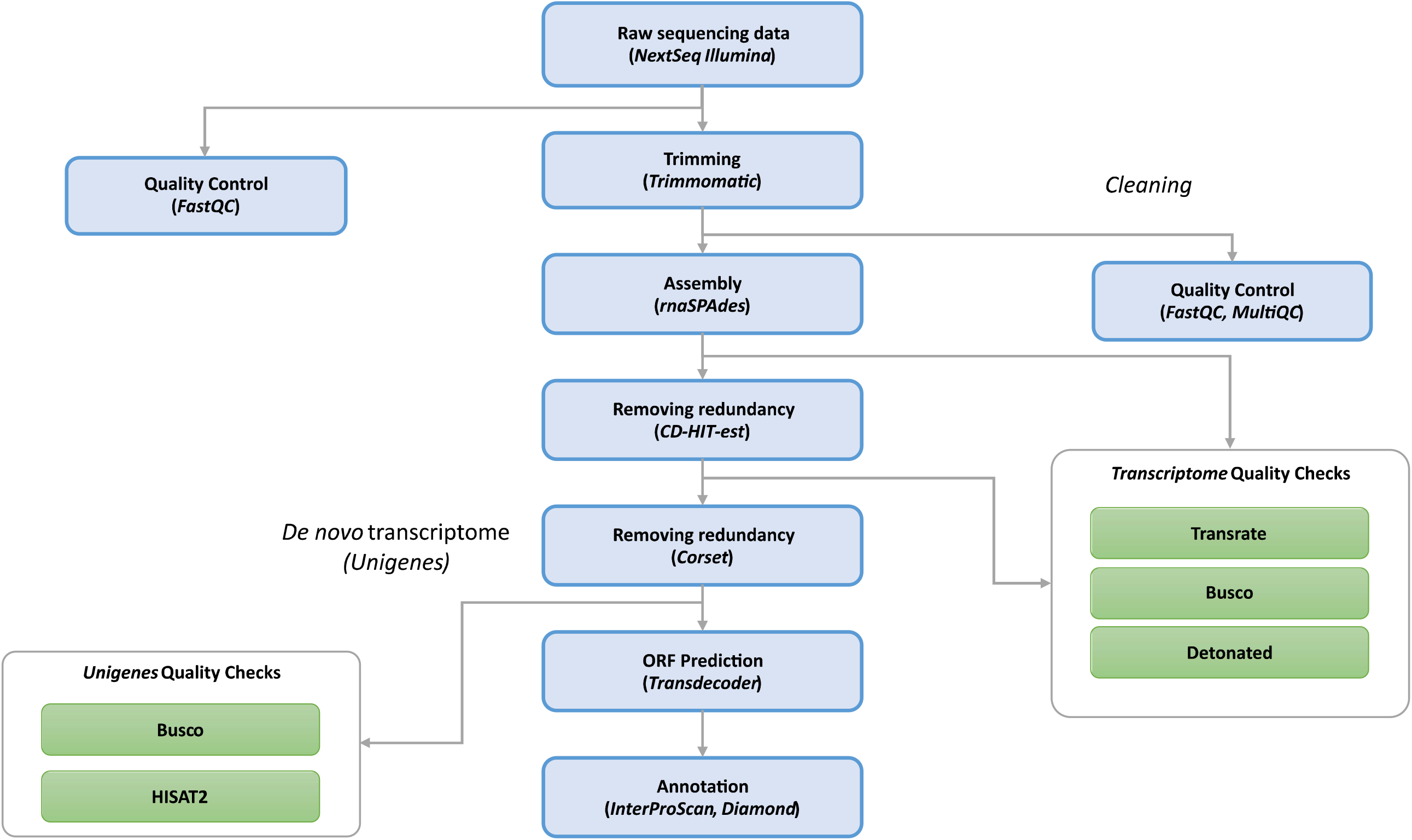
Workflow of the bioinformatic pipeline, from raw input data to annotated contigs, for the *de novo* transcriptome assembly of *B. pachypus.*

To construct an optimized *de novo* transcriptome, avoiding chimeric transcripts, we employed rnaSPAdes^23^, a tool for *de novo* transcriptome assembly from RNA-Seq data implemented in the SPAdes v.3.14.1 package. rnaSPAdes automatically detected two k-mer sizes, approximately one third and half of the maximal read length (the two detected k-mer sizes were 45 and 67 nucleotides, respectively). At this stage, a total of 1,118,671 assembled transcripts were generated by rnaSPAdes runs, with an average length of 689.41 bp and an N50 of 1474 bp (Table 2).

**Table 2.** Similarity rate of newly assembled transcripts versus the *de novo* transcriptome of *B. pachypus*.

|  | After rnaSPAdes | After CD-HIT-est | After Corset (Unigenes) |
| --- | --- | --- | --- |
| **Basic parameters** |  |  |  |
| Total transcripts | 1118671 | 896992 | 267959 |
| Mean transcripts length (bp) | 689.41 | 616.32 | 799 |
| N50 | 1474 | 1082 | 2314 |
| GC content (%) | 40.15 | 39.93 | 39.87 |
| **Transrate v.1.0.3** |  |  |  |
| Transrate Assembly Score | 0.0563 | 0.1281 | - |
| Transrate Optimal Score | 0.0877 | 0.178 | - |
| Transrate Optimal Cutoff | 0.0138 | 0.0138 | - |
| good contigs | 991122 | 822552 | - |
| p good contigs | 0.89 | 0.92 | - |
| **Busco v.3.0.2** |  |  |  |
| Complete Buscos (C) | 245 (96.1%) | 240 (94.1%) | 246 (96.5 %) |
| Complete and single-copy Buscos (S) | 157 (61.1%) | 186 (72.9 %) | 157 (61.6 %) |
| Complete and duplicated Buscos (D) | 88 (34.5 %) | 54 (21.2 %) | 89 (25.3 %) |
| Fragmented Buscos (F) | 4 (1.6 %) | 8 (3.1 %) | 8 (3.1 %) |
| Missing Buscos (M) | 6 (2.3 %) | 7 (2.8 %) | 1 (0.4 %) |
| Total Busco groups searched | 255 | 255 | 255 |
| **Detonate v.1.9** |  |  |  |
| Score | -28610853057.11 | -31067445817.60 | - |
| BIC_penalty | -10912136.97 | -11301742.60 | - |
| Prior_score_on_contig_lengths_(f_function_canceled) | -2403141.71 | -2100226.61 | - |
| Prior_score_on_contig_sequences | -1072761315.55 | -993903654.87 | - |
| Data_likelihood_in_log_space_with_out_correction | -27529522562.34 | -30064989468.40 | - |
| Correction_term_(f_function_canceled) | -4746099.45 | -4849274.88 | - |

Results from the assembly procedures were validated through three independent validator algorithms implemented in BUSCO^24^ v.4.1.4, DETONATE^25^ v.1.11 and TransRate^26^ v.1.0.3. These tools generate several metrics used as a guide to evaluate error sources in the assembly process and provide evidence about the quality of the assembled transcriptome. Busco provides a quantitative measure of transcriptome quality and completeness, based on evolutionarily-informed expectations of gene content from the near-universal, ultra-conserved eukaryotic proteins (eukaryota_odb9) database. Detonate (DE novo TranscriptOme rNa-seq Assembly with or without the Truth Evaluation) is a reference-free evaluation method based on a novel probabilistic model that depends only on the assembly and the RNA-Seq reads used to construct it. Transrate generates standard metrics and remapping statistics. No reference protein sequences were used for the assessment with Transrate. The main metrics resulted from the assembly validators are shown in Table 2 *(“Before CD-HIT-est”* column). After this triple assessment validation step, the result of the assembly procedure become the input for the CD-HIT-est v.4.8.1^27^ program, a hierarchical clustering tool used to avoid redundant transcripts and fragmented assemblies common in the process of *de novo* assembly, providing unique genes. CD-HIT-est was run using the default parameters, corresponding to a similarity of 95%. Subsequently, a second validation step was launched on the CD-HIT-est output file. To refine the final transcriptome dataset, a further hierarchical clustering step was performed by running CORSET v1.06^28^. Then, the output of CORSET was validated by BUSCO, and quality assessment was performed with HISAT2^29,30^ by mapping the trimmed reads to the reference transcriptome (unigenes). Results from all validation steps are shown in Table 2 and discussed in the “Technical Validation” paragraph.

Finally, the CORSET output was run on TransDecoder^31,32^, the current standard tool that identifies long open read frames (ORFs) in assembled transcripts, using default parameters. TransDecoder by default performs ORF prediction on both strands of assembled transcripts regardless of the sequenced library. It also ranks ORFs based on their completeness, and determines if the 5 ‘end is incomplete by looking for any length of AA codons upstream of a start codon (M) without a stop codon. We adopted the “Longest ORF” rule and selected the highest 5 AUG (relative to the inframe stop codon) as the translation start site.

### Transcriptome annotation

We employed different kinds of annotations for the de novo assembly. We introduced DIAMOND^33^, an open-source algorithm based on double indexing that is 20,000 times faster than BLASTX on short reads and has a similar degree of sensitivity. Like BLASTX, DIAMOND attempts to determine exhaustively all significant alignments for a given query. Most sequence comparison programs, including BLASTX, follow the seed-and-extend paradigm. In this two-phase approach, users search first for matches of seeds (short stretches of the query sequence) in the reference database, and this is followed by an ‘extend’ phase that aims to compute a full alignment. The following parameter settings were applied: DIAMOND-fast DIAMOND BLASTX-t 48 -k 250 -min-score 40; DIAMOND-sensitive: DIAMOND BLASTX -t 48 -k 250 -sensitive -min-score 40.

Contigs were aligned with DIAMOND on Nr, SwissProt and TrEMBL to retrieve the corresponding best annotations. An annotation matrix was then generated by selecting the best hit for each database. Following the analysis of BLASTX against Nr, SwissProt and TremBL, we obtained respectively: 123,086 (64.57%), 77,736 (40.78%), 122,907 (64.48%) contigs. The results obtained following the analysis with BLASTP against Nr, SwissProt and TrEMBL were 96,321 (50.53%), 57,877 (30.36%) and 97,256 (51.02%) contigs respectively. All the information on the resulting datasets is resumed in Table 3.

**Table 3.** Summary of homology annotation hits on the different databases queried in this study.

| Number of homology annotated contigs on different databases |  |  |
| --- | --- | --- |
| Database | Number of BLASTX results | Number of BLASTP results |
| Nr | 123086 (64.57 %) | 96321 (50.53 %) |
| SwissProt | 77736 (40.78 %) | 57877 (30.36 %) |
| TrEMBL | 122907 (64.48 %) | 97256 (51.02 %) |

The output obtained by the BLASTX annotation consisted in a total of 77391 sequences simultaneously mapped on the three queried databases (i.e., Nr, SwissProt and TrEMBL). The output obtained following the BLASTP annotation consisted in a total of 57704 sequences simultaneously mapped on the three databases. Venn diagrams are presented in Figure 3, showing the redundancy of the annotations in the different databases for both DIAMOND BLASTX (Figure 3a) and DIAMOND BLASTP (Figure 3b). Furthermore, the ten most represented species and the ten hits of the gene product obtained respectively with BLASTX and BLASTP by mapping the transcripts against the reference database Nr are shown in Figure 4 and 5. Since BLASTX translated nucleotide sequence searches against protein sequences the BLASTX results are more exhaustive than BLASTP results. Contigs were also processed with InterProScan^34^ to detect InterProScan signatures. The InterPro database (http://www.ebi.ac.uk/interpro/) integrates together predictive models or ‘signatures’ representing protein domains, families and functional sites from multiple, diverse source databases: Gene3D, PANTHER, Pfam, PIRSF, PRINTS, ProDom, PROSITE, SMART, SUPERFAMILY and TIGRFAMs. The obtained InterProScan results for all the unigenes are available on Figshare in the form of Tab Separated Values (tsv) file format, which includes the GO and KEGG annotated contigs, respectively.

**Figure 3.**
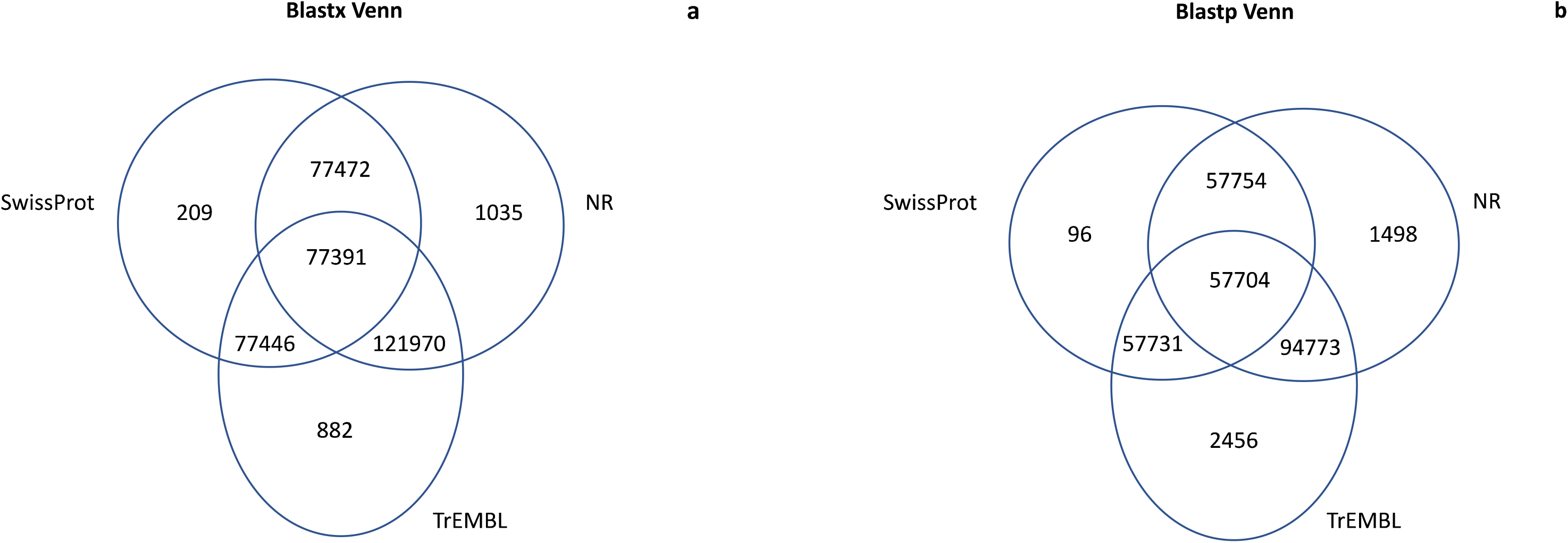
Venn diagrams for the number of contigs annotated with DIAMOND (BLASTX (a) and BLASTP (b) functions) against the three databases: Nr, SwissProt, TREMBL.

**Figure 4.**
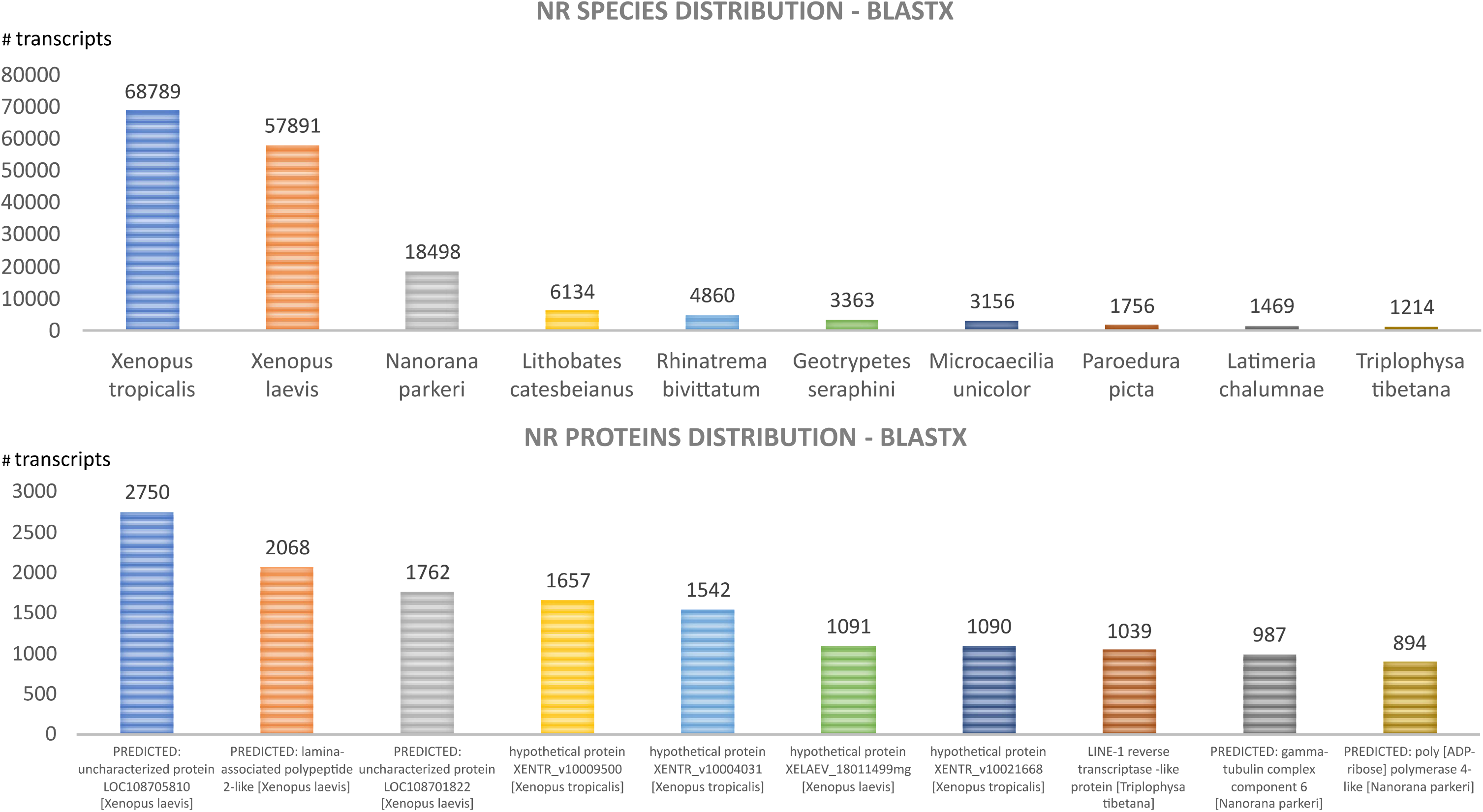
Most represented species and gene product hits. Top 10 best species (a) and protein (b) hits present in the reference database (Nr, BLASTX).

**Figure 5.**
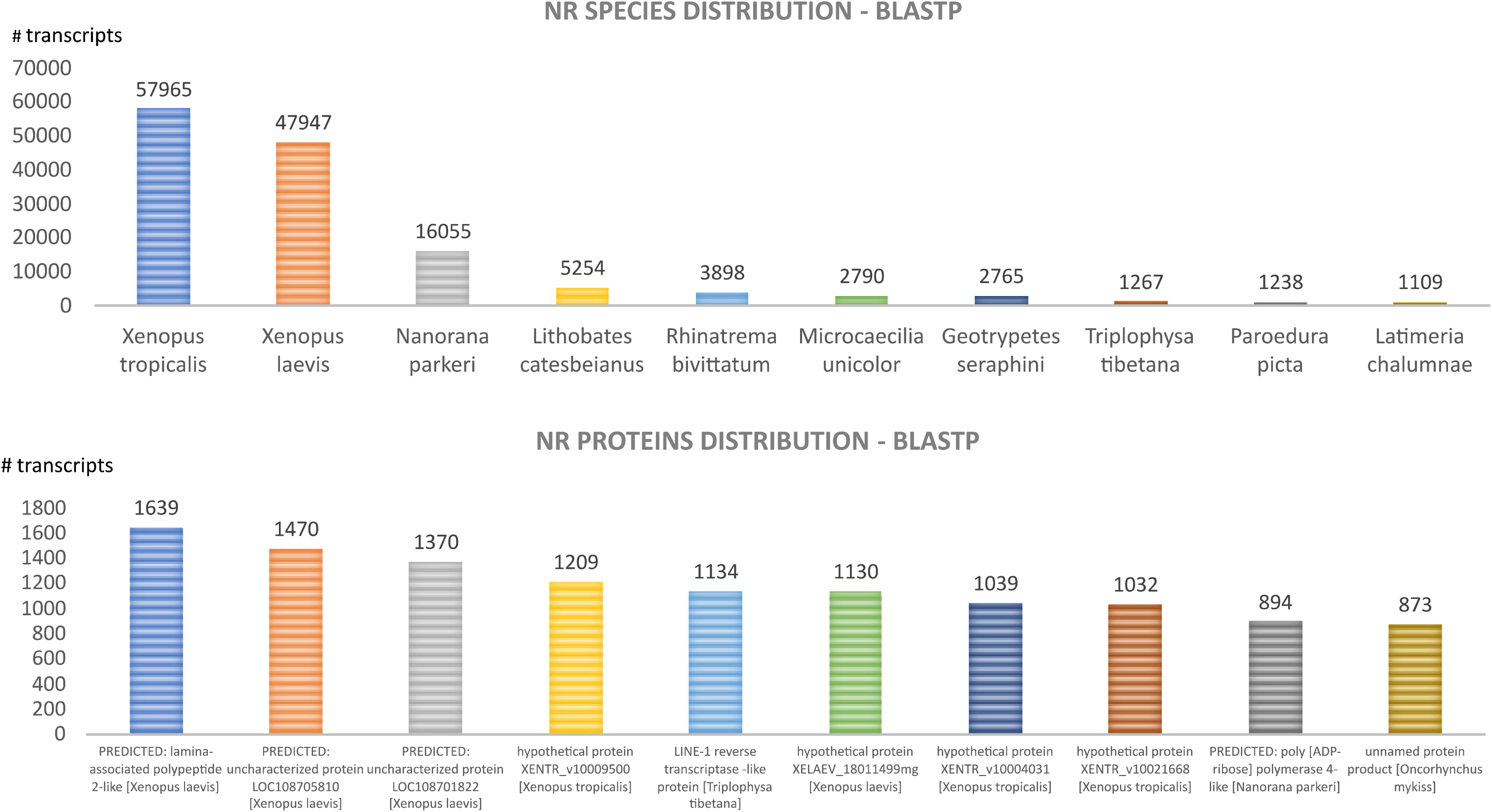
Most represented species and gene product hits. Top 10 best species (a) and protein (b) hits present in the reference database (Nr, BLASTP).

### Comparison with *Bombina orientalis* brain transcriptome

We compared the brain *de novo* transcriptome of *B. pachypus* with the brain *de novo* transcriptome of *B. orientalis,* recently produced in the frame of a prey-catching conditioning experiment^17,18^. The *B. orientalis* transcriptome resource was downloaded from GEO archive of NCBI (https://www.ncbi.nlm.nih.gov/geo/query/acc.cgi?acc=GSE171766). To make the datasets comparable, we first performed ORF prediction on *B. orientalis* trascriptome using Transdecoder, using default settings. Results from the *B. orientalis* trascriptome ORF prediction are available in Figshare at the following link https://doi.org/10.6084/m9.figshare.20319633/). We also applied the *makedb* function implemented in DIAMOND to create the protein database index. Then, we aligned the *B. pachypus* predicted coding sequences and proteins (query files) against the *B. orientalis* protein *database* (reference) using DIAMOND BLASTX and BLASTP, respectively. We obtained 167041 matches from BLASTX and 156248 matches for BLASTP. Results from the BLASTX and BLASTP comparisons, and the most matched proteins, are available on Figshare (link available in next paragraph).

### Data Records

We generated six files corresponding to the RNA-seq samples of the brain tissue of the six *B. pachypus* individuals analyzed for this study. The six files were deposited in the NCBI Sequence Read Archive database, under project identification number PRJNA764013; the NCBI accessions for each individual are listed in Table 1 (Run ID SRR15927729 - SRR15927734). The datasets containing the rnaSPAdes transcriptome assembly, post CD-Hit-Est assembly, post corset assembly (unigenes), predicted ORF and homology and functional annotation files were deposited in Figshare (Project description: https://doi.org/10.6084/m9.figshare.c.5696179; Assembly: https://doi.org/10.6084/m9.figshare.16945270; Annotation: https://doi.org/10.6084/m9.figshare.16945264; comparison with *Bombina orientalis* transcriptome: https://doi.org/10.6084/m9.figshare.20319633).

### Technical Validation

#### Quality of the raw reads and assembly validation

To assess overall data quality, we performed quality checks using FastQC and MultiQC for all samples before and after adaptor/sequence trimming. The mean read counts per quality score were higher than 35 (Figure 1a). The mean quality scores in each base position were higher than 35 (Figure 1b). The mean sequence lengths were 126-130 bp (Figure 1c). The mean per sequence GC content was 40% (Figure 1d).

Transcriptome assembly validation was done using Busco, Detonate and Transrate. Results from the triple validation step are shown in Table 2, and contain the scores obtained from the execution of the three analysis tools, both before and after running CD-HIT-est.

Using HISAT2^30^ (a fast and sensitive alignment program for mapping next-generation sequencing reads, DNA and RNA), we verified that more than 91% of the reads were mapped back to the assembled transcriptome of the *B. pachypus* thus indicating a proper quality sequence reconstruction. Figure 5 shows the number of raw reads, paired-reads after trimming, and trimmed paired-reads that are mapped against B. pachypus de novo transcriptome.

**Figure 6.**
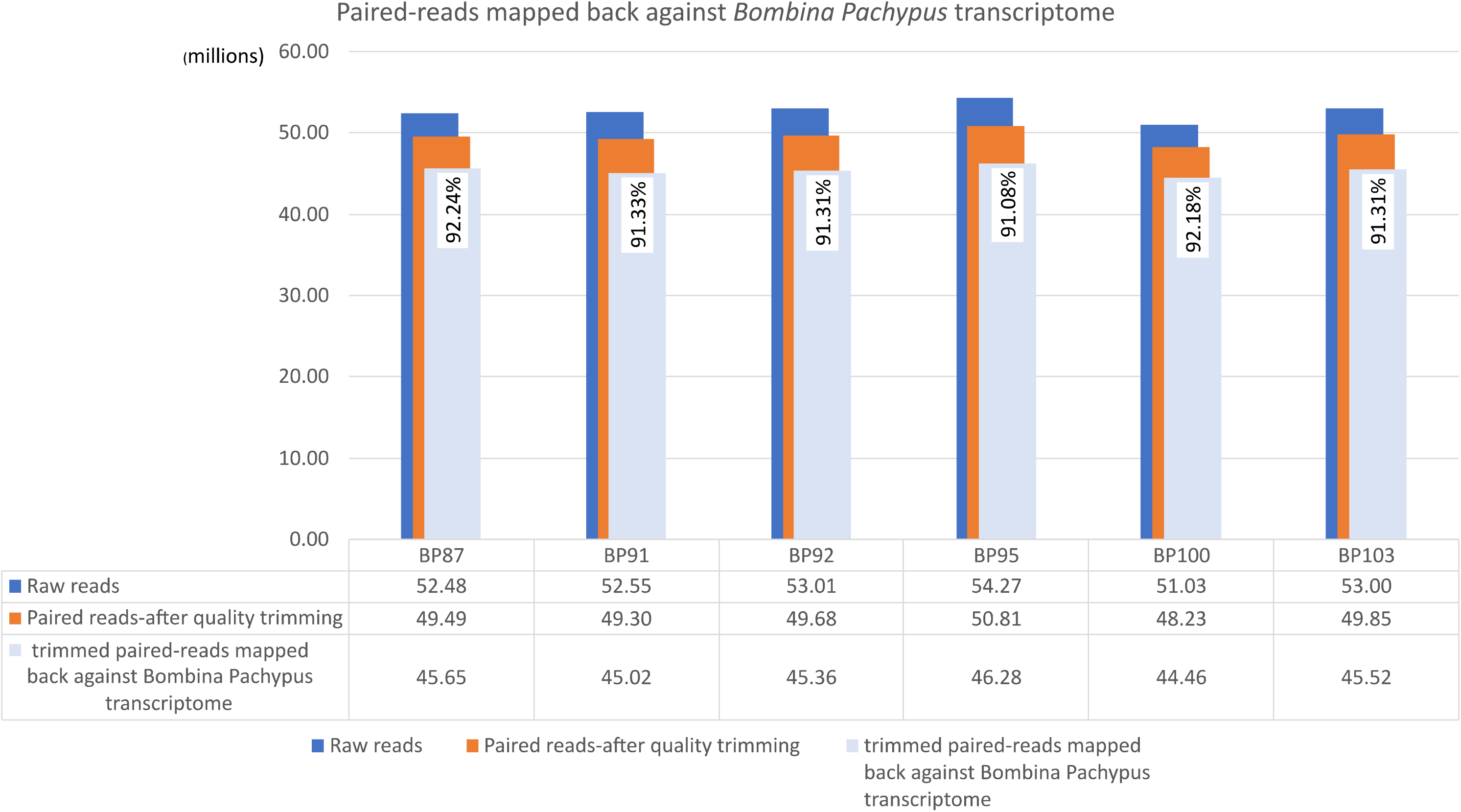
For each sample we have in blue the representation of total paired-reads, in orange the total paired-reads after the adapter removal and quality trimming and in azure we have the trimmed paired-reads mapped mapped-back against the *B. pachypus* assembled *de novo* transcriptome.

Transrate assessment showed increased values for the “Transrate Optimal Score” item following hierarchical clustering using CD-HIT-est, passing from 0.088 to 0.178, and for the “Transrate Assembly Score” item, passing from 0.056 to 0.128 (more than twice). The value of “good contigs” increased after CD-HIT-est (due to redundancy removal), but with a value of 0.92 for the final assembly. Transrate also reported a value of GC around 40 % after each validation step. The transcriptome obtained after CD-HIT-est included a total of 896,992 transcripts with a mean transcript length of 616.32 bp and an N50 of 1082 bp, with a value above the 94% of completeness for Busco assessment. In terms of redundancy removal the further step of CORSET clustering produced a real improvement. In fact, the final version of the assembled transcriptome included 267,959 transcripts with a mean transcript length of 799 bp, the N50 value equals to 2314 and a value above the 96% for Busco assessment, improving the previous results computed by the CD-HIT-est tool. As shown in Table 2, CORSET greatly improved the assembled transcriptome removing redundancy and reducing the number of transcripts, thus improving the quality scores of the final assembly.

#### Quality control of annotation

The transcriptome was functionally annotated by performing DIAMOND and InterProScan. By selecting the best hit for Nr, SwissProt and TrEMBL databases, the annotation matrix generated with DIAMOND has led to the results listed in table 3. In particular 77,391 (BLASTX) and 57,704 (BLASTP) contigs were annotated in all the three databases, NR, Swissprot, Trembl.

InterProScan provided as result the corresponding InterPro accession numbers and, among other accession IDs, the GO and Kegg annotation. It produced a total of 32142 annotated contigs, being 4747 contigs GO-annotated and 1025 contigs KEGG-annotated.

## Code availability

All the software programs used in this article (*de novo* transcriptome assembly, pre and post-assembly steps, and transcriptome annotation) are listed in the Methods paragraph. In case of no details on parameters, the programs were used with the default settings.

## Acknowledgments

We are grateful to Michela Paoletti for her support during the laboratory procedures and to Jessica Di Martino for her work on the transcriptome annotation. We acknowledge the CINECA for the availability of high-performance computing resources and the ELIXIR-ITA HPC@CINECA initiative for providing HPC resources to our projects: 1) name of the call “Call ELIXIR-ITA CINECA (2020– 2021)”, P.I. Andrea Chiocchio, name of the project “ELIX4_chiocchi”; 2) name of the call “Call ELIXIR-ITA CINECA (2021–2022)”, P.I. Tiziana Castrignanò, name of the project “ELIX4_castrign2”. This study was supported by grants from the Italian Ministry for Education, University and Research (Prin project: 2017KLZ3MA), and from the Aspromonte National Park.

## Author contributions

DC conceived and financed the study; AC e DC designed the experiment; AC, RB and GM performed sample collection and preparation; AC coordinated the RNA extraction and sequencing; TC designed and coordinated the bioinformatic analysis; PL and TC performed reads quality assessment, reads alignment on transcriptome, transcriptome annotation and validation; AC, PL and TC wrote the manuscript; DC, TC, AC, RB, PL and GM reviewed the manuscript.

## Competing interests

The Authors declare no competing interests.

## Notes

### Competing Interest Statement

The authors have declared no competing interest.

## References

1. Carere, C. & Maestripieri, D. Animal Personalities: Behavior, Physiology, and Evolution. (Chicago: University of Chicago Press, 2013).

2. Jensen, P. Behaviour epigenetics–the connection between environment, stress and welfare. Appl. Anim. Behav. Sci. 157, 1–7 (2014).

3. Van Oers, K., de Jong, G., van Noordwijk, A. J., Kempenaers, B. & Drent P. J. Contribution of genetics to the study of animal personalities: a review of case studies. Behaviour 142, 1185–120610 (2005).

4. Van Oers, K. & Sinn, D. L. The quantitative and molecular genetics of animal personality. In: Carere, C. & Maestripieri, D. editors. Animal Personalities: Behavior, Physiology, and Evolution. (Chicago: University of Chicago Press; p. 148–200, 2013).

5. Ellegren, H. Genome sequencing and population genomics in non-model organisms. Trends Ecol. Evol. 29, 51–63 (2014).

6. Umbers, K. D. L., Lehtonen, J., & Mappes, J. Deimatic displays. Curr. Biol. 25, R58eR59, (2015).

7. Joron, M., & Mallet, J. L. Diversity in mimicry: paradox or paradigm?. Trends Ecol. Evol. 13, 461–466, (1998).

8. Arenas, L. M., & Stevens, M. Diversity in warning coloration is easily recognized by avian predators. J. Evol. Biol. 30, 1288–1302, (2017).

9. Richards-Zawacki, C. L., Yeager, J., & Bart, H. P. No evidence for differential survival or predation between sympatric color morphs of an aposematic poison frog. Evol. Ecol. 27, 783–795 (2013).

10. Rönkä, K. Evolution of signal diversity: predator-prey interactions and the maintenance of warning color polymorphism in the wood tiger moth *Arctia plantaginis*. Jyväskylä studies in biological and environmental science 339, (2017).

11. Lawrence, J. P. et al. Weak warning signals can persist in the absence of gene flow. Proc. Natl. Acad. Sci. USA 116, 19037–19045, (2019).

12. Chiocchio, A., Martino, G., Bisconti, R., Carere, C., Canestrelli D. Shock or jump: deimatic behavior is repeatable and polymorphic in a yellow-bellied toad. bioRxiv 2022.04.29.489992; doi: https://doi.org/10.1101/2022.04.29.489992 (2022).

13. Koolhaas, J.M., de Boer, S.F., Coppens, C.M., Buwalda, B. Neuroendocrinology of coping styles: towards understanding the biology of individual variation. Front Neuroendocrinol. 31(3), 307–21 (2010).

14. Whitfield C. W., Cziko, A. M., & Robinson, G. E. Gene expression profiles in the brain predict behavior in individual honey bees. Science 302, 296–299, (2003).

15. Rey, S., Boltana, S., Vargas, R., Roher, N., & Mackenzie, S. Combining animal personalities with transcriptomics resolves individual variation within a wild-type zebrafish population and identifies underpinning molecular differences in brain function. Mol. Ecol. 22, 6100–15, (2013).

16. Bell, A. M., Bukhari, S. A., & Sanogoc, Y. O. Natural variation in brain gene expression profiles of aggressive and nonaggressive individual sticklebacks. Behavior 153, 1723–1743, (2016).

17. Lewis, V., Laberge, F., Heyland, A. Temporal Profile of Brain Gene Expression After Prey Catching Conditioning in an Anuran Amphibian. Front Neurosci 3, 1407 (2020).

18. Lewis V, Laberge F, Heyland A. Transcriptomic signature of extinction learning in the brain of the fire-bellied toad, Bombina orientalis. Neurobiol Learn Mem. 184, 107502 (2021)

19. Harris, R. M., & Hofmann, H. A. Neurogenomics of behavioral plasticity. In Landry, C. R. & Aubin-Horth N. editors. Ecological genomics. (Springer Science, pp. 149–168, 2014).

20. Ewels, P., Magnusson, M., Lundin, S. & Kaller, M. MultiQC: summarize analysis results for multiple tools and samples in a single report. Bioinformatics 32, 3047–8 (2016).

21. Bolger, A. M., Lohse, M. & Usadel, B. Trimmomatic: a flexible trimmer for Illumina sequence data. Bioinformatics 30, 2114–20 (2014).

22. Castrignanò, T., Gioiosa, S., Flati, T., Cestari, M., Picardi, E., Chiara, M., Fratelli, M., Amente, S., Cirilli, M., Tangaro, M.A., Chillemi, G., Pesole, G., Zambelli, F. ELIXIR-IT HPC@CINECA: high performance computing resources for the bioinformatics community. BMC Bioinformatics. 21(Suppl 10), 352 (2020).

23. Bushmanova, E., Antipov, D., Lapidus, A., Prjibelski, A.D. rnaSPAdes: a *de novo* transcriptome assembler and its application to RNA-Seq data. Gigascience 8, giz100 (2019).

24. Simão, F. A., Waterhouse, R. M., Ioannidis, P., Kriventseva, E. V. & Zdobnov, E. M. Busco: Assessing genome assembly and annotation completeness with single-copy orthologs. Bioinformatics 31, 3210–3212 (2015).

25. Li, B. et al. Evaluation of *de novo* transcriptome assemblies from RNA-Seq data. Genome Biol. 15, 1–21 (2014).

26. Smith-Unna, R., Boursnell, C., Patro, R., Hibberd, J. M. & Kelly, S. Transrate: Reference-free quality assessment of *de novo* transcriptome assemblies. Genome Res. 26, 1134–1144 (2016).

27. Fu, L., Niu, B., Zhu, Z., Wu, S. & Li, W. CD-HIT: Accelerated for clustering the next-generation sequencing data. Bioinformatics 28, 3150–3152 (2012).

28. Davidson, N.M., Oshlack, A. Corset: enabling differential gene expression analysis for de novo assembled transcriptomes. Genome Biol. 15(7), 410 (2014)

29. Kim, D., Langmead, B. & Salzberg, S. L. HISAT: a fast spliced aligner with low memory requirements. Nat. Methods. 12, 357–360 (2015).

30. Pertea, M., Kim, D., Pertea, G., Leek, J. T. & Salzberg, S. L. Transcript-level expression analysis of RNA-seq experiments with HISAT, StringTie and Ballgown. Nat. Protoc. 11, 1650–67 (2016).

31. Signal, B., & Kahlke, T. Borf: Improved ORF prediction in *de novo* assembled transcriptome annotation. Preprint at https://www.biorxiv.org/content/10.1101/2021.04.12.439551v1([A-Z]) (2021).

32. Tang, S., Lomsadze, A., Borodovsky, M. Identification of protein coding regions in RNA transcripts. Nucleic Acids Res. 43, (2015).

33. Buchfink, B., Xie, C. & Huson, D. Fast and sensitive protein alignment using DIAMOND. Nat. Methods. 12, 59–60 (2015).

34. Hunter, S. et al. InterPro: the integrative protein signature database. Nucleic Acids Res. 37, D211–5.

